# New solid-state optical pH sensors for cell analysis

**DOI:** 10.64898/2026.08.04.742867

**Authors:** Liang Li

**Affiliations:** School of Biochemistry and Cell Biology, University College Cork, Cork, Ireland

**Keywords:** Optochemical pH sensors, Fluorescent Porphyrins, Extracellular acidification rate, Cell metabolism and bioenergetics, Sensor cytotoxicity

## Abstract

Monitoring pH and extracellular acidification rate (ECA) in biological samples containing live mammalian cells can provide valuable information on the glycolytic activity and bioenergetic status of cells. Compared to pH electrodes, optochemical pH sensors look more advantageous, since they allow rapid, non-invasive parallel analysis of multiple samples with stable readout of pH. We have developed new fluorescent pH sensors based on hydrophobic protonable metal-free porphyrins,OEP and OEPK, embedded in a plasticized PVC matrix containing a proton transfer agent. These pH sensors provide internally-referenced calibration-free operation, both in ratiometric intensity and lifetime-based detection modes. Sensor development included optimization of the indicator dye and its photophysical characteristics, screening of different proton transfer agents to minimize sensor toxicity, tuning of the protonation range and pKa, long-term storage stability and response time studies. Optimised pH sensor coatings were then deposited on plastic substrates (96-well microplates) and used for real-time monitoring of Extracellular Acidification Rate (ECAR) for cultured cancer cells and 3D spheroid structures on standard laboratory equipment (multi-label plate reader and confocal FLIM microscope). The advanced pH sensors tailored for use with biological samples have high potential for cell analysis and related applications.

## 1. INTRODUCTION

pH is a measure of acidity or basicity of an aqueous solution, one of the most frequently analyzed parameters. Physiological pH values normally range between 6.0 and 8.0, spanning down to pH 4-6 for some specialized cellular compartments)[1]. Determination of pH values in biological fluids, cells and tissue samples is important for medicine, biotechnology, tumor radiotherapy, corrosion monitoring, food and chemical sciences [2-4]. The gold standard of pH analysis is glass pH electrode, which can provide rapid, stable, and reliable readout in a wide pH range, which is proportional to sample pH. However, due to its significant size (>1 mm, even if miniaturized), susceptibility to electrical interferences, difficulty of sterilization and rigid design, glass electrodes are poorly suitable for biological samples containing cells [2, 5]. Thus, new approaches to pH monitoring in solutions are in high demand.

Optochemical pH sensors have emerged as an alternative to the electrode. They allow simple, non-destructive measurement of multiple samples, and localized pH gradients. They are well suited for in vitro and in vivo measurements and show high flexibility, robustness, accuracy, biocompatibility and multiplexing capabilities with other dyes/sensors [6, 7]. It is also possible to develop pH sensors with internal referencing, which allow for calibration-free operation, demonstrate high specificity, brightness, reversible and rapid response to pH changes, tunable dynamic range. They can also operate in different detection modalities and on available instruments[8].

We present a new family of solid-state optochemical pH sensors based on fluorescent porphyrin dyes OEP and OEPK, incorporated in a proton-permeable plasticized PVC matrix containing a proton transfer agent (PTA). The sensors have adjustable pKa values in the physiological range (6.0 ≤ pH ≤ 8.0), low cytotoxicity, and can be used both in the internal referenced ratiometric intensity and fluorescence lifetime modes. They were deposited on common plastic substrates (microwell plates), and demonstrated in the measurement of extracellular acidification rate (ECAR) in biological samples containing cells.

## 2. METHODS

### Methods

OEP and OEPK dyes were initially incorporated in PVC matrices/coatings plasticized with dioctyl sebacate (DOS), and containing tetrachlorophenyl borate (TCPB) as a proton transfer agent. Later on, TCPB was replaced with sodium tetrakis[3,5-bis(trifluoromethyl)phenyl]borate (Na-TFMB), which shows better solubility in plasticized PVC and exerts low toxicity on cells. Sensor coatings were produced by preparing a precursor cocktail (2% PVC and 4% DOS, OEP/OEPK and Na-TFMB borate in THF/chloroform), and spotting it on a polyester film (Mylar, Du Pont) or into wells of round bottom 96-well polypropylene microplate, followed by drying at room temperature overnight. Fluorescence spectra were measured on Cary Eclipse spectrofluorometer (Agilent, USA). Ratiometric intensity ECAR analysis was performed using Victor2 reader (Perkin-Elmer, Waltham, MA, USA) a prompt fluorescence mode. Fluorescence lifetime based analysis of ECAR was performed on a confocal TCSPC-FLIM microscope (Becker & Hickl, Berlin, Germany). HCT116 cells were cultured in McCoy’s 5A medium supplemented with 10% foetal bovine albumin.

## 3. RESULTS

The Ocaethylporphine (OEP) and ocaethylporphine-ketone (OEPK) dyes are both excitable at 400 nm (Soret band) and emit at >600 nm, with sharp well-resolved bands of their neutral and protonated forms (Figure 1). This allows OEP and OEPK based solid-state pH sensors to be used in a ratiometric intensity mode on standard fluorescence readers.

**Figure 1.**
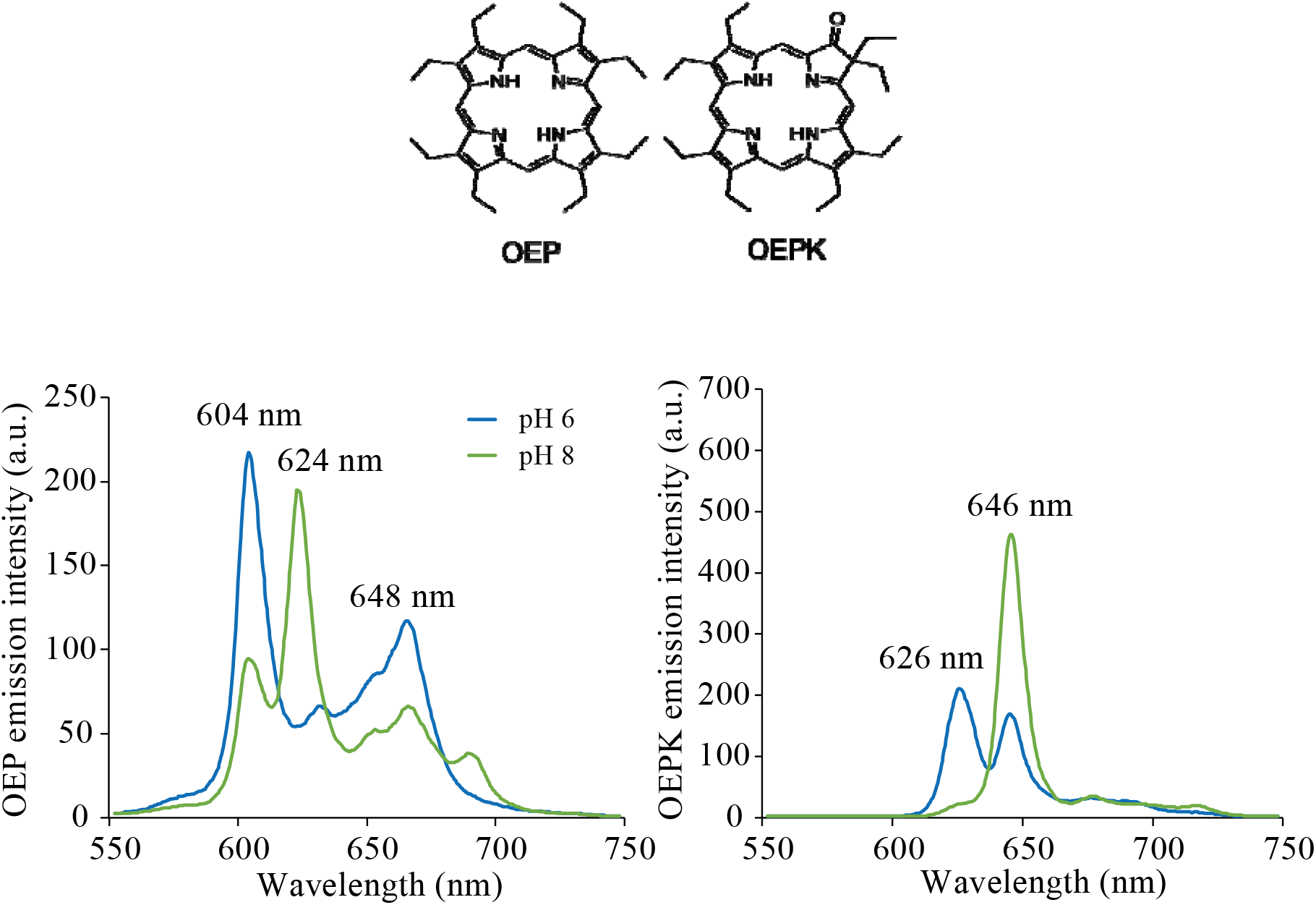
Chemical structures (A) and emission spectra (B) of OEP and OEPK at pH 6.0 (blue line) and pH 8.0 (green line).

We also found that protonation/deprotonation is accompanied by reversible changes in fluorescence lifetime (FLT) of the dyes (Figure 2). The magnitude of FLT response was much greater than the existing lifetime-based pH sensors. This allows OEP- and OEPK-based sensors to be used on existing FLT detectors e.g., confocal fluorescence lifetime imaging (FLIM) microscope with time-correlated single photon counting (TCSPC) detector.

**Figure 2.**
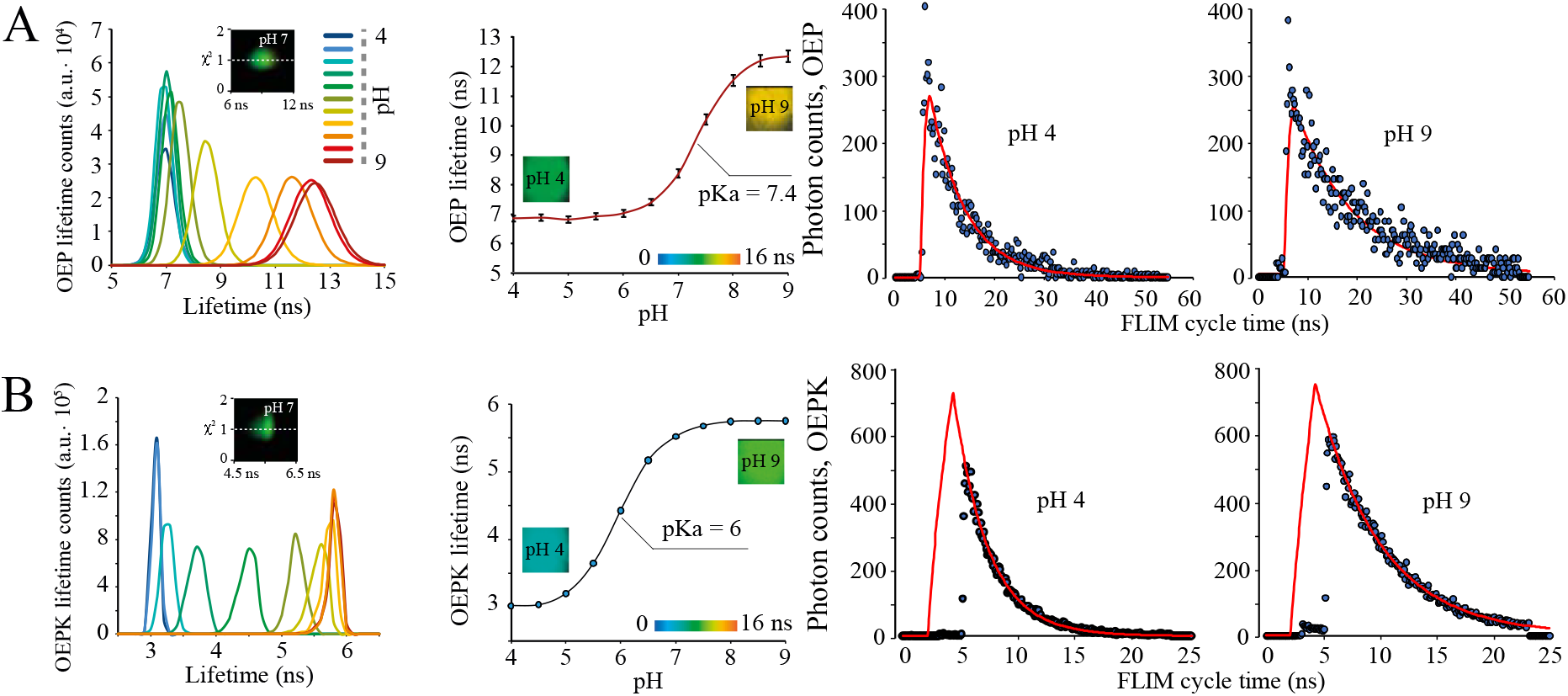
Analysis of pH responses of the OEP and OEPK based sensors by FLIM confocal microscopy. A-B. Calibrations curves and decay traces of OEP and OEPK sensors at different pH, generated on the confocal FLIM microscopy.

OEP and OEPK dyes were previously used in pH sensors based on plasticized PVC matrix and lipophilic borate salts (TCPB) as proton transfer agent (PTA) [9]. However, these sensors were not tested with biological samples containing live respiring cells. We found that main drawbacks of these sensors are: the prominent impact of TCPB on cell viability, and low solubility of TCPB in cocktail solution, leading to dye aggregation and inhomogeneous sensor coatings.

To reduce the cytotoxicity of TCPB and optimize the sensor coatings, we evaluated the effects of washing, protection with collagen IV or hydrogel D4 over-layer on sensor toxicity (Figure 3A-B). Overnight incubation of the sensor in PBS did not reduce its toxicity, while overnight incubation in RPMI medium with 5% FBS significantly reduced toxicity and vanished sensor response to pH. The over-layer of D4 hydrogel (5% in 90% ethanol or 3.75% D4 in 67.5% ethanol) was seen to increase sensor signals, reduce toxicity, but also eliminated pH sensitivity. The collagen IV over-layer kept the pH sensitivity but did not improve toxicity.

**Figure 3.**
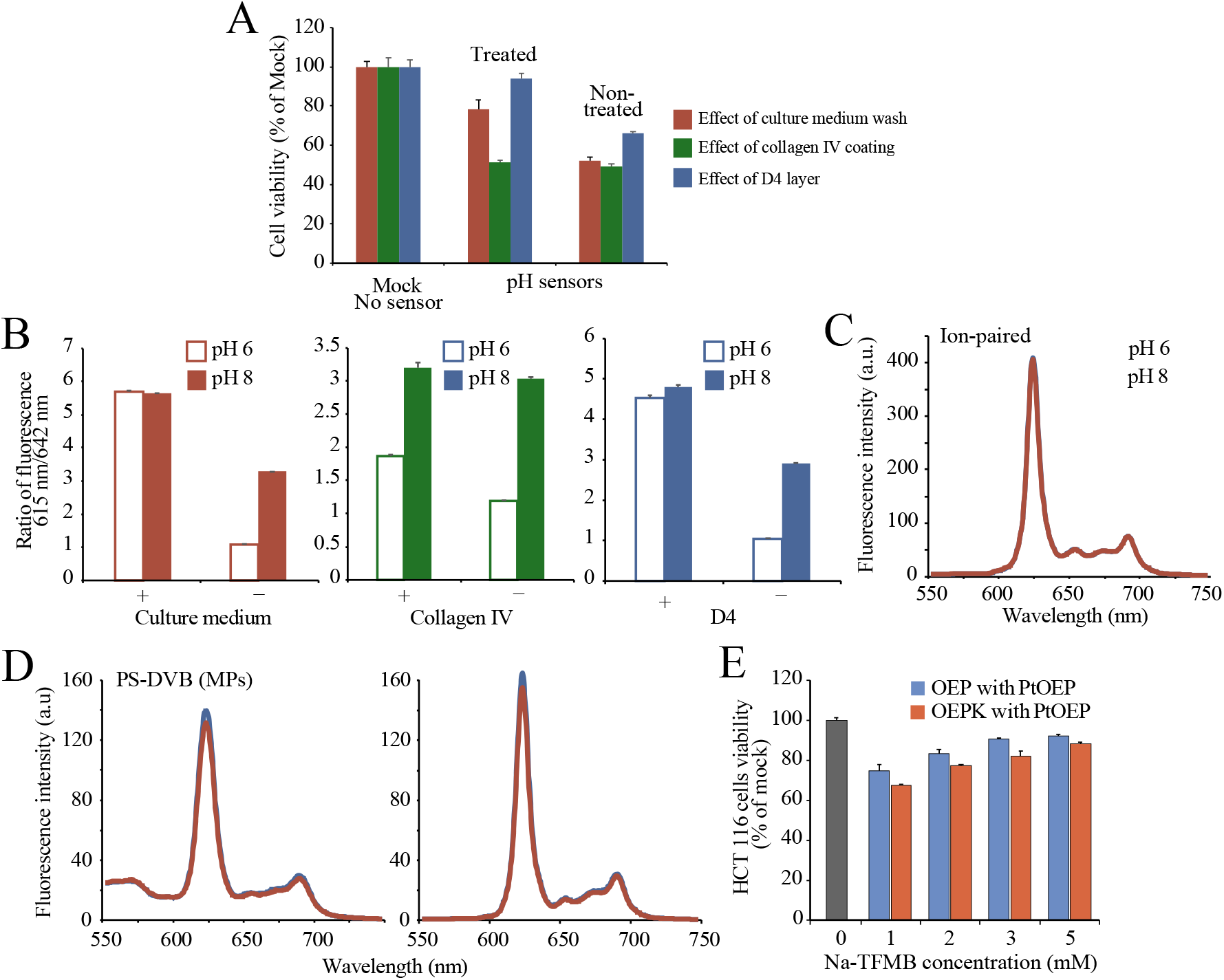
Optimization of the sensor toxicity. A. Cell viability tests with PC12 (effects of medium prewash and collagen) and HCT116 cells (effect of D4 protective layer). B. Effects of sensor pre-treatments on the 615/642 nm ratio. C. Effect of OEP+-borate-ion pairs on sensor pH response. D. Effect of PS-DVB Microparticles on sensor pH sensitivity (3 mM and 30 mM TCPB). E. The effect of Na-TFMB concentrations on HCT 116 cell viability.

We also attempted to develop pH sensors based on OEP+-borate-ion pairs and PS-DVB microparticles (MPs) to reduce toxicity (Fig3C-D). OEP+-borate-ion pairs were produced in CH_2_Cl_2_-HCl solvent system, followed by the incorporation of these particulate sensors in 5% D4 (EtOH solvent) or 6% plasticized PVC (THF). The D4 and plasticized PVC sensor coatings on Mylar showed practically no pH response (spectral corresponded to unprotonated OEP). When the PS-DVB MPs were co-impregnated with OEP and TCPB using the same molar ratio as in the sensor (0.25 mM and 3 mM), they showed no spectral response to pH, both in aqueous suspension and in D4 coatings. Maximizing the TCPB content (30 mM) still did not allow protonation and the appearance of a characteristic peak at 603 nm.

Although these special steps partly reduced the cytotoxicity of TCPB based sensors, they also decreased or even eliminated the response to pH. Therefore, they were ruled out. To solve the problem of sensor toxicity, we screened a small panel of alternative PTAs, which included three borate (Na-TFMB, Li-PFPB, and Na-TPB) and two organic phosphates salts (diphenyl phosphate and 1-1’-binapthyl-2,2’-diyl hiylhydrogen-phosphate). Form this panel, the pH sensors based on Na-TFMB (PTAs) showed low cytotoxicity and excellent pH-sensing performance with tunable pKa values (Fig 3E, Fig.5A).

Prior to the experiments with biological samples, the Mylar film with sensor coatings were prepared (200 µL sensor cocktail on each 22×22 mm strips) by spin-coating, inserted diagonally in a quartz cuvette and measured on Cary Eclipse in kinetic mode to assess their response time to pH changes. The results showed that at low spinning speed, sensor response time was slower but intensity signals were higher (Fig.4). Optimal spinning speed was determined as 1000 rpm. Under these settings, sensor pH response on Mylar film was fully reversible and complete in ∼2.5 mins. We also studied response time of sensor coatings deposited on the 96-wells microplate, measuring the fluorescence ratio signal on a Victor2 reader. Surprisingly, the pH response in microplate was fully reversible and even faster ∼1.5 mins, and sensor signal was more stable (Fig 5B).

**Figure 4.**
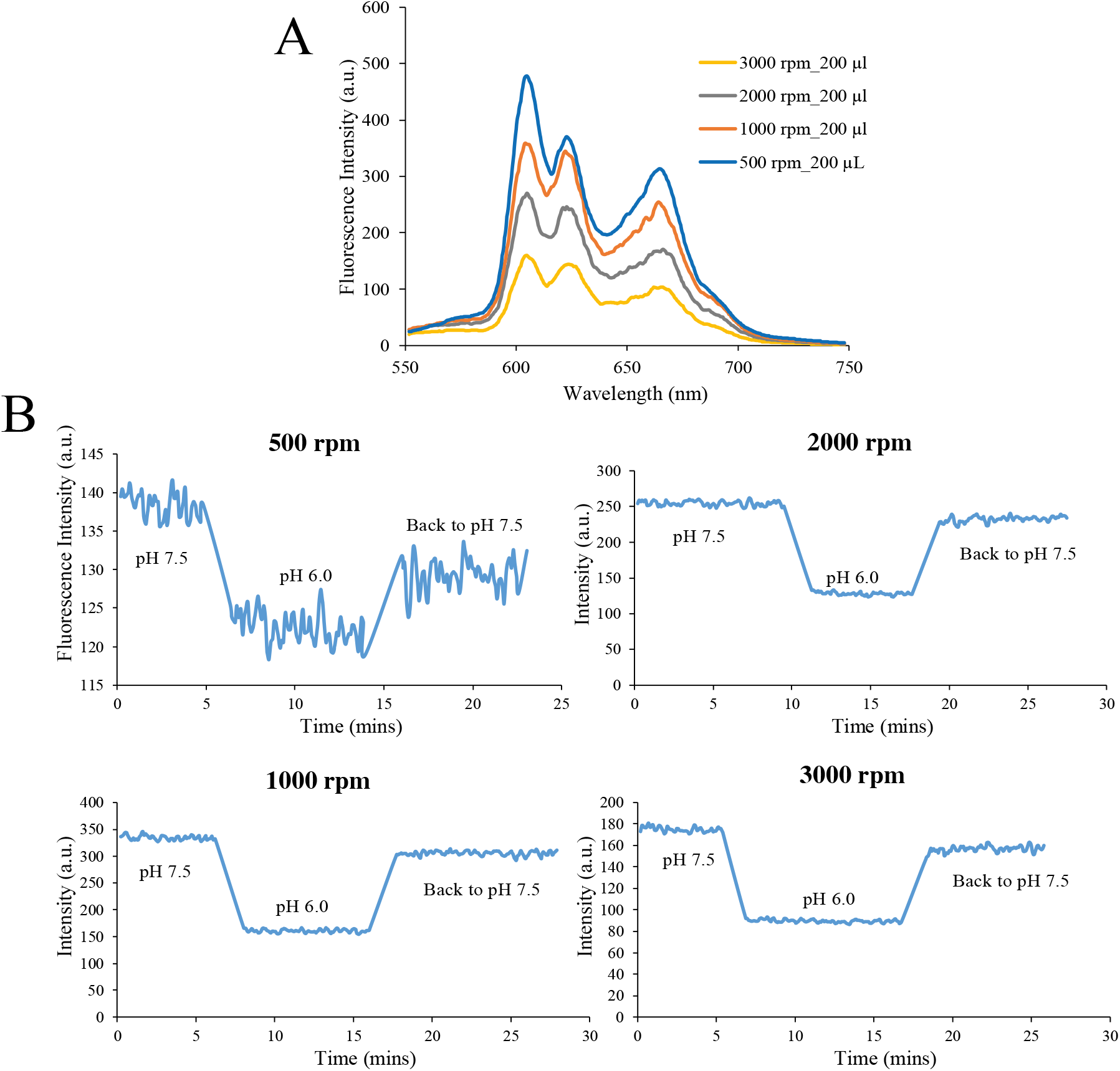
Optimization of sensor time response on Cary Eclipse. A. Fluorescence spectrums of sensor coatings (OEP) on Mylar films produced at different spin speeds by a Spin Coater. B. Time response of OEP sensor coatings on Mylar films tested in a kinetic mode.

**Figure 5:**
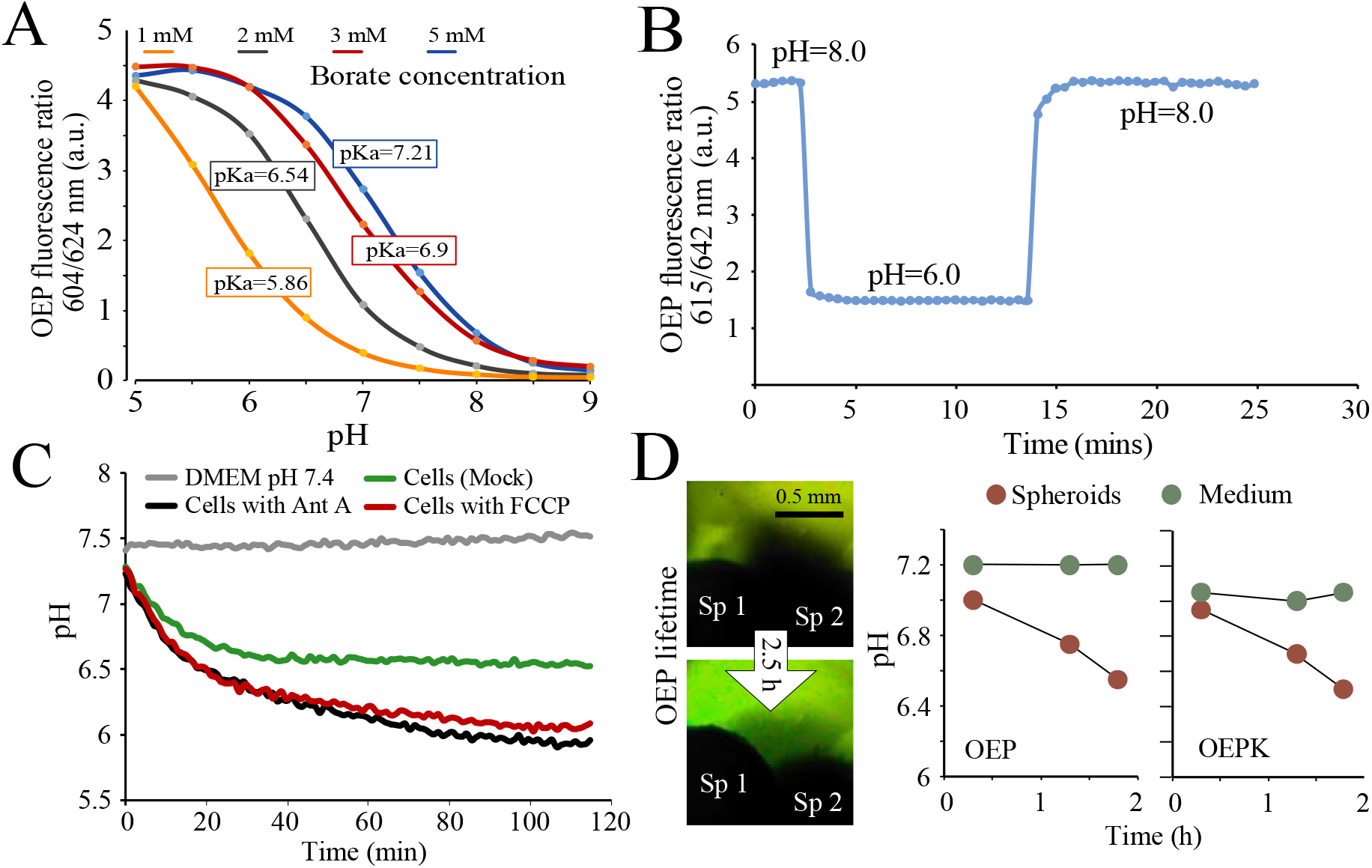
Biological applications of pH sensors. A. Tunable pKa values based on different Na-TFMB concentrations on Cary Eclipse. B. Response time for sensor signals upon pH transmission on victor2 plate reader. C-D. Extracellular acidification rate (ECAR) analysis for HCT 116 cells by a Victor2 plate reader (C) and its 3D spheroids by a FLIM confocal microscope (D).

Then the optimized sensors were applied in biological samples for the extracellular acidification rate (ECA) analysis. Sensor coatings were deposited on a polypropylene 96-well microplates, and HCT 116 cells treated with FCCP, and Ant A (effector compound) or DMSO (as mock, negative control) were measured and assessed. The fluorescence intensity ratio and lifetime (ns) signals, measured on a Victor2 plate reader and FLIM confocal microscope, were converted into pH values, using the corresponding calibration curves and equations, The strong pH response was observed for the samples with cells (Fig5C-D). Compared to mock, there was a larger decrease in pH for FCCP and even larger for Ant A treatment. These results report on the activated glycolysis and lactate release upon the inhibition of oxidative phosphorylation [10]. The 3D spheroid structures made of HCT 116 cells were analyzed on a confocal FLIM microscope for their ECA. The OEP and OEPK sensors both showed the anticipated pH responses, and their results agreed with each other.

Furthermore, the optimized OEP and OEPK based sensor also showed excellent stability over 12 months of storage (in dry form, at 4oC), with stable pKa values and no significant degradations of the sensor dye or coating (Fig 6).

**Figure 6.**
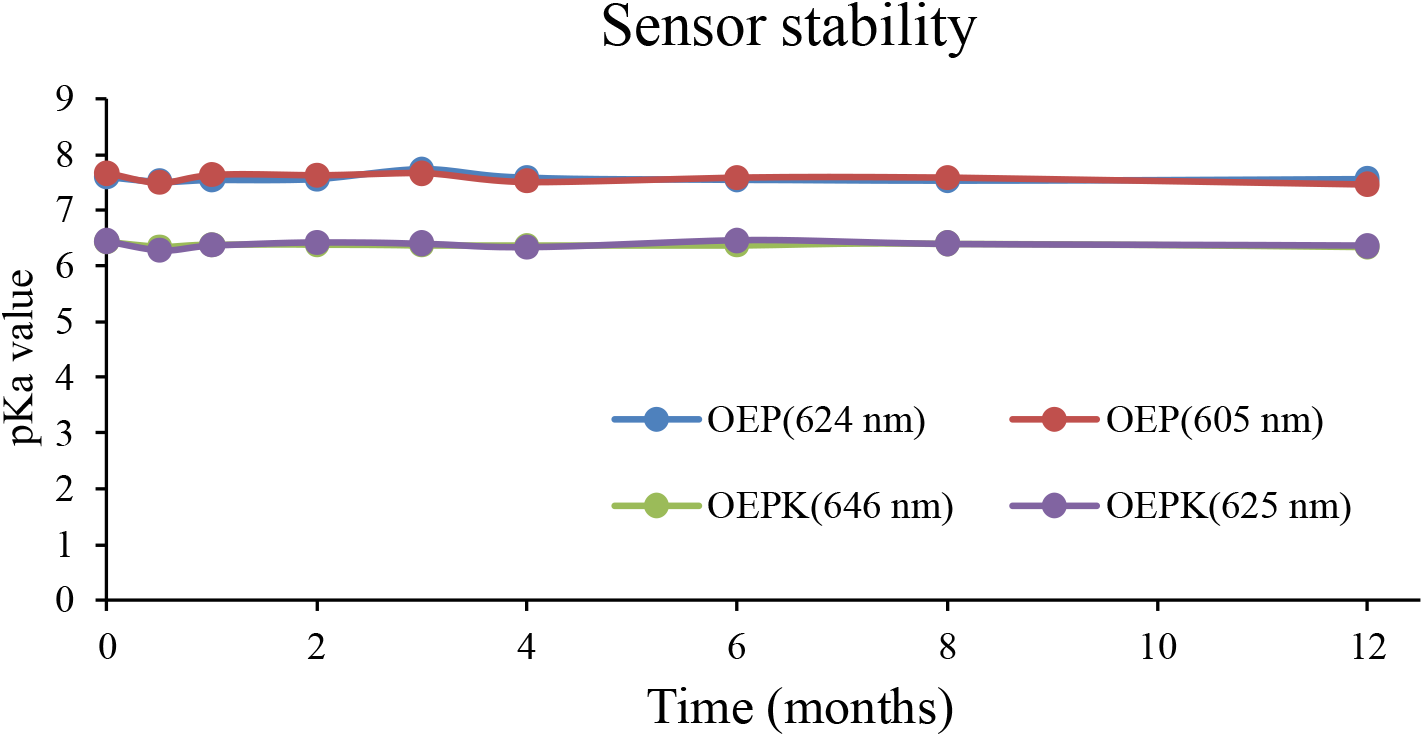
The storage stability of OEP and OEPK based sensor. The pKa values were calculated based on protonation and deprotonation peaks of OEP and OEPK on Cary Eclipse (N=3).

## Notes

### Competing Interest Statement

The authors have declared no competing interest.

